# SEARCH_16S: A new algorithm for identifying 16S ribosomal RNA genes in contigs and chromosomes

**DOI:** 10.1101/124131

**Authors:** Robert C. Edgar

## Abstract

SEARCH_16S is a new algorithm that annotates 16S ribosomal RNA genes in microbial genomes and metagenomic sequences. Word counting is used to identify candidate segments, then conserved motifs are used to identify homologous loci close to the gene boundaries. SEARCH_16S has >99% sensitivity to known 16S genes.

## Introduction

Microbial 16S ribosomal RNA is the most widely sequenced gene in biology, currently accounting for 14 million sequences in Genbank (Dec. 2016). Despite its importance, annotations of the 16S gene in microbial genomes are unreliable because commonly-used methods such as BLAST (Altschul *et al*., 1997) and HMMer (Eddy, 2009) are not accurate for this purpose (Freyhult *et al*., 2007; Lagesen *et al*., 2007). A specialized tool is therefore desirable. To the best of my knowledge, only one such tool has been published, namely RNAmmer (Lagesen *et al*., 2007), which I was unable to obtain. I therefore developed SEARCH_16S, a new method for finding 16S genes. SEARCH_16S identifies segments with a high frequency of 13-mers in known 16S genes, then searches within each such segment for conserved motifs close to the beginning and end of the gene. Finding a pair of motifs within the expected length range confirms the presence of the gene and provides consistent, homologous endpoints. It would be preferable to identify the true endpoints of the functional sequence, but the 16S gene is spliced out of the ribosomal operon by mechanisms that are not fully understood and lacks known sequence signals analogous to start and stop codons for protein-coding genes (Shajani *et al*., 2011). I validated SEARCH_16S on finished prokaryotic genomes and curated SSU databases, finding that it has >99% sensitivity to known genes and no unambiguous false positives in control datasets containing metazoan sequences and random sequences.

## Methods

### Signature words

SEARCH_16S uses the set of all 13-mers (*signature words*) found in GG97, the Greengenes database (DeSantis *et al*., 2006) v13.5 clustered at 97% identity. The algorithm starts by searching for regions with high frequencies of these words. The word length of 13 was determined as follows. Shorter words give better sensitivity to novel genes because more of them are conserved between related sequences, while longer words give better specificity because they are less likely to be found by chance in unrelated sequences. Almost all possible 8-mers (98%) are found in GG97, while only 12% of possible 13-mers and 2% of 15-mers are present (Fig. 1). I therefore chose length 13 as a compromise between optimizing sensitivity and specificity. Given that 12% of all possible 13-mers are present in GG97, a background frequency of ∼12% is expected in non-16S sequence and an elevated frequency »12% is expected where there is high sequence similarity to GG97.

**Fig 1.**
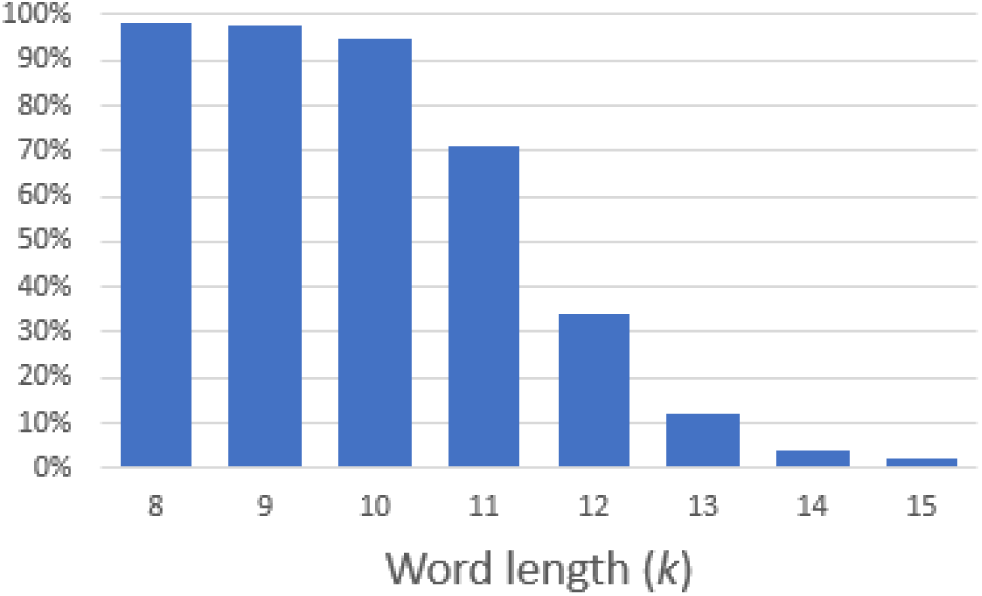
Word occurrence in GG97 as a function of length. For each word length (*k*) in the range 8 to 15 the number of distinct words found in GG97 is shown as a fraction 4*^k^*, the number of distinct possible words. For *k*<12, most possible words are found in the database and these values of *k* are therefore not suitable for signature words that distinguish 16S from other sequences.

### Candidate segments

To find regions with elevated frequencies, SEARCH_16S calculates the number of signature words (*density*) for every window of length 1,000bp. In a window of this length, the expected background density is ∼120 because there is a ∼12% chance that a given 13-mer is present in GG97. In a known gene, the density should be close to 1,000, and in a novel gene the density should be much higher than the background (Fig. 2). A region with elevated frequency of signature words (a *candidate segment*) is identified as a maximal series of consecutive positions having density ≥500, with margins of half the window length (500bp) added at each end. The margins are needed because with a window of length *w*, there will be *w*/2 positions at the beginning of the gene whose windows contain at least one word outside of the gene, giving lower densities at those positions, and similarly there will be *w*/2 windows which contain at least one position after the end. Thus, for a gene of length 1,500bp which is present in GG97, we expect to find 500bp at the start of the gene where the density rises from 120 to 1,000, a flat peak of 500bp in the middle of the gene where the density is maximum, then 500bp at the end of the gene where the density falls from 1,000 back to 120, as seen in the lower panel in Fig. 2. An additional margin of 200bp is added at both ends to allow for lower sequence identity, which reduces the number of positions with density above the detection threshold of 500. The total margin is thus 700bp, and if 500bp consecutive positions have density ≥500 the candidate segment will then have length 500 + 2×700 = 1,900bp.

**Fig. 2.**
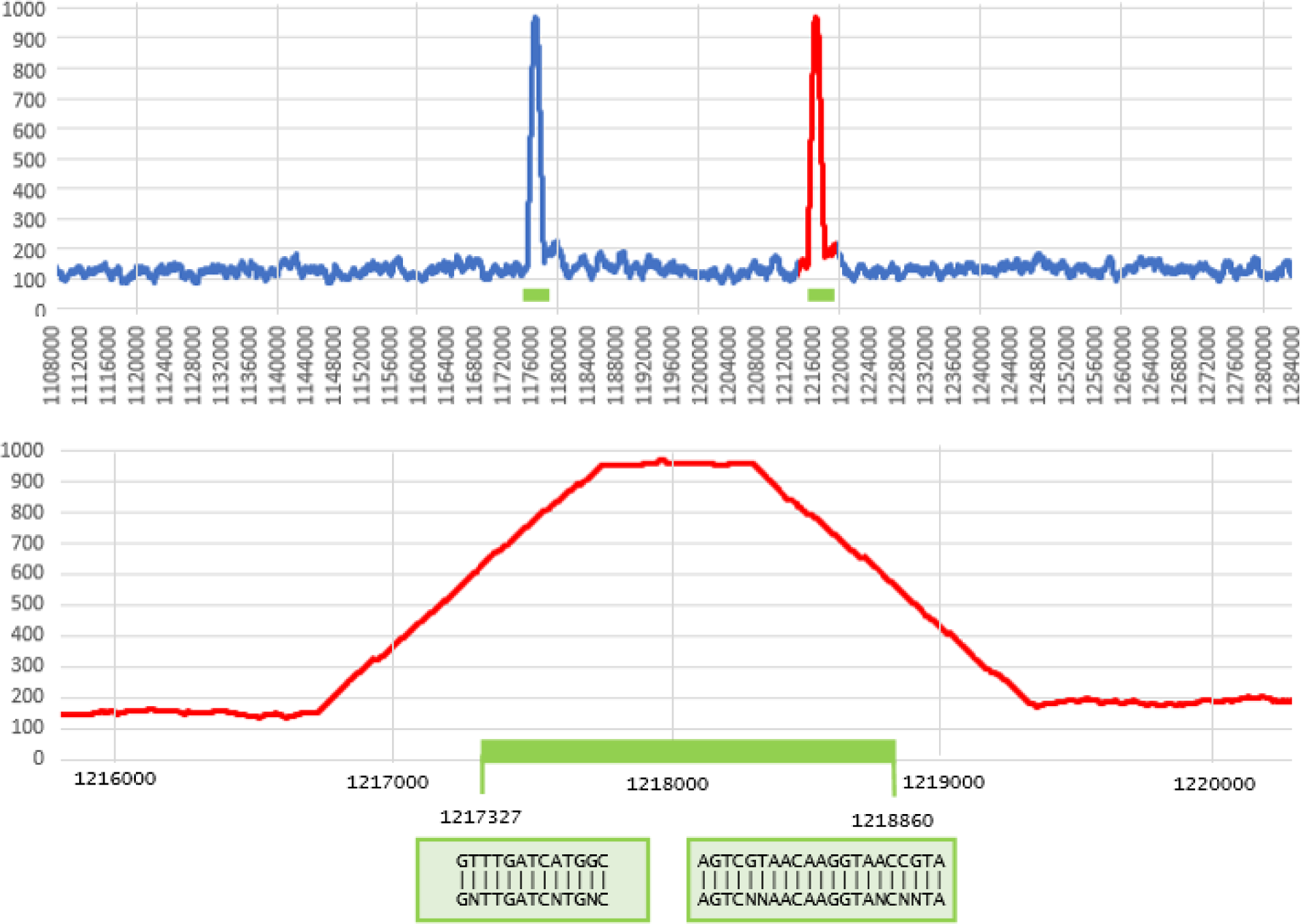
Signature word density for a region of the *E. coli* chromosome reverse strand. In the top panel, the density of signature 13-mers over windows of length 1,000bp is shown for positions 1,108,000 – 1,284,000 in Genbank sequence AP009048.1. Most positions have a density close to the expected background of ∼120 words per window. The two 16S genes in this region (green bars) are visible as spikes where the density approaches 1,000. The lower panel shows the region from positions 1,216,000 to 1,220,000 where the second gene is located. The trapezoidal shape of the density is explained by windows which contain some words before/ after the beginning / end of the gene; the flat peak of length approx. 500bp is due to windows that contain only 16S words. The boundary motifs are found at positions 1,217,327 (C11F) and 1,218,860 (C1512R).

### Boundary motifs

The start and end of the gene are found by searching a candidate segment for the *boundary motifs* C11F = GNTTGATCNTGNC and C1512R = AGTCNNAACAAGGTANCNNTA, allowing up to four mismatches and choosing the match with fewest differences. If matches to both motifs are found, and the sequence truncated to the motifs is between 1,000bp and 2,500bp in length, then that sequence is reported as a 16S gene. Otherwise, the segment is reported as a *fragment*. A fragment may be a valid 16S gene in which one or both motifs are missing (e.g. in a contig containing a partial gene), a valid 16S gene in which the motifs are not recognized because the sequence is too far diverged, or a homologous ribosomal gene such as 18S. The alignments of C11F and C1512R to the *E. coli* 16S sequence (Genbank J01859.1) are shown in Fig. 3. I use the convention that motifs are named *Cn*F if they are close to the start of the gene or *Cm*R if they are close to the end of the gene, where *n* is the position in J01859.1 to which the *first* base of the motif aligns and *m* is the position in J01859.1 where the *last* base of the motif aligns. All motif sequences are given on the forward strand. Thus, SEARCH_16S annotates bases 11 through 1512 of J01859.1 as 16S for a total length of 1,501 bases, omitting 10 bases at the true start of the gene and 29 bases at the end.

**Fig 3.**
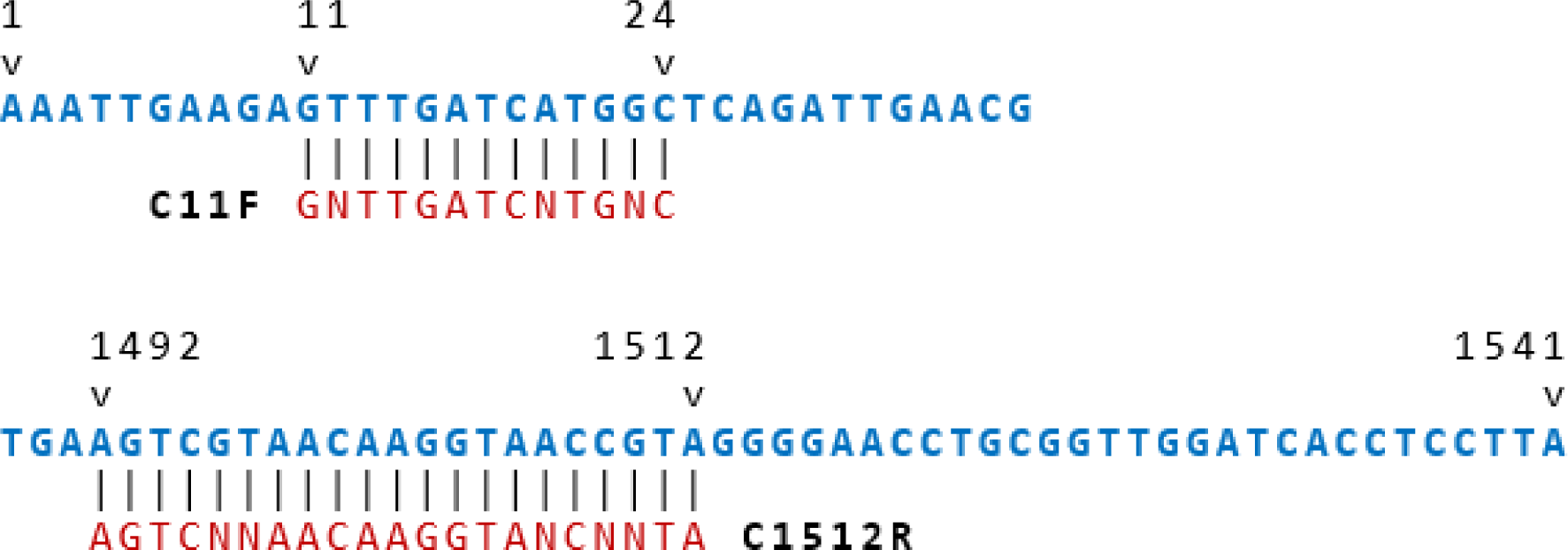
Alignment of C11F and C1512R motifs to the E. coli 16S sequence (Genbank J01859.1). The gene is 1,541bp. The alignment of the C11F motifs starts at position 11 and the C1512R motif ends at position 1512. These motifs are used to determine the start and end of an annotation, causing 10 and 29 bases to be omitted from the start and end of the *E. coli* gene, respectively. The omitted regions include loop secondary structures with less sequence conservation, presumably because they are not constrained by base pairing.

**Fig. 4.**
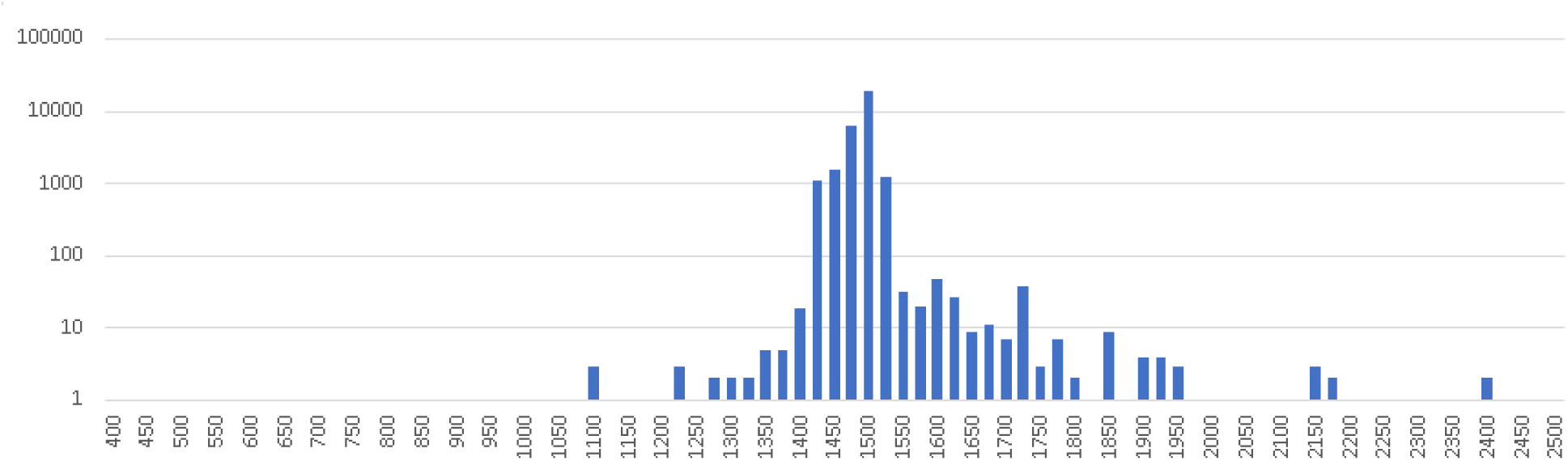
Length distribution of genes reported by SEARCH_16S in finished genomes. The histogram shows lengths binned into intervals of 25nt. The peak is at the 1500-1549nt bin which contains 18904 / 29450 (64%) of the genes, with only 24 (0.08%) having lengths <1400nt. Genes were clustered at 97% identity to reduce redundancy caused by species such as *E. coli* that have several assemblies.

## Results

### Validation datasets

To test the sensitivity of SEARCH_16S I used all finished prokaryotic genomes in Genbank and three large curated databases containing small subunit (SSU) ribosomal sequences: SILVA (Pruesse *et al*., 2007) v23, RDP (Maidak *et al*., 2001) downloaded 30th Sept. 2016 and Greengenes v13.5. I identified the set of complete prokaryotic genomes in assembly_summary_genbank.txt (downloaded from NCBI on 20th Dec. 2016) and downloaded the sequences and feature tables for those assemblies. SILVA contains Archaea, Bacteria and Eukaryote subsets, RDP contains Archaea, Bacteria and Fungi, and Greengenes contains only Bacteria and Archaea. To investigate specificity in reporting 16S but not other SSU genes, I used the eukaryotic subset of the SILVA SSU database and the metazoan subset of the NCBI nt (non-redundant nucleotide) database (Sayers *et al*., 2012). Almost all metazoa are eukaryotes whose small subunit genes are conventionally classified as 18S rather than 16S. I also created a large dataset of random sequences as a further test of specificity. Results on these tests are summarized in Table 1.

**Table 1.** 16S hits in the test datasets. Here, a hit is a gene or fragment reported by SEARCH_16S. For finished genomes, the *Genes* column reports the number of genomes that have at least one reported gene, not the total number of genes reported in those genomes. The datasets are “Complete” genomes (see main text), Greengenes (all sequences), RDP (prokaryotic subset), SILVA (prokaryotic subset), SILVA (eukaryotic subset), Metazoa, the subset of the NCBI nt database classified as metazoan, and Random (100k sequences of length 1M). Most of the sequences in SILVA, RDP and SILVA are truncated (Fig. 6), explaining the large number of reported fragments.

| Dataset | Sequences | Genes | Fragments | No hit | Hit% |
| --- | --- | --- | --- | --- | --- |
| Finished genomes | 6,487 | 6,475 | 9 | 12 | 99.95% |
| Greengenes | 1,262,986 | 87,593 | 1,175,393 | 0 | 100.00% |
| RDP (prok.) | 3,356,808 | 126,318 | 2,708,298 | 522,192 | 84.44% |
| SILVA (prok.) | 1,636,081 | 168,037 | 1,467,876 | 168 | 99.99% |
| SILVA (euk.) | 120,702 | 5 | 74 | 120,623 | 0.07% |
| Metazoa | 12,379,862 | 25 | 1,268 | 12,378,581 | 0.01% |
| Random | 100,000 | 0 | 0 | 100,000 | 0.0% |

### Complete prokaryotic genomes

I tested the 6,487 assemblies in Genbank that were annotated as prokaryotic (classified as Bacteria or Archaea by the NCBI taxonomy) and as a “Complete Genome”. SEARCH_16S reported at least one 16S gene in 6,475 (99.8%) of these genomes. The twelve assemblies with no reported 16S gene are summarized in Table 2. I used BLAST (Altschul *et al*., 1997) to search these assemblies against GG97 to check for significant local alignments. BLAST found no hits with E-values <10 in the three assemblies where SEARCH_16S reported neither genes nor fragments, indicating that their small-subunit rRNA genes are missing and these assemblies are not in fact “complete”. SEARCH_16S reported fragments in five assemblies which had high-identity BLAST hits (97% to 100%) covering only partial sequences in GG97, strongly suggesting assembly errors. SEARCH_16S reported fragments in four assemblies where BLAST hits had high identities (99% to 100%) and were long enough (1,340nt to 2,201nt) to be consistent with full-length genes. Thus, the results on this test can be interpreted as indicating 100% sensitivity of SEARCH_16S if (a) fragments are regarded as possible genes that require further analysis, and (b) it is assumed that if one gene in a genome is successfully found, then all genes in that genome will be found. The latter assumption is reasonable because 16S paralogs are usually identical or have high sequence identity (Acinas *et al*., 2004). If the four reported fragments that may be intact genes are uncharitably considered to be false negatives, then the sensitivity of SEARCH_16S on this test is 99.94%.

**Table 2.** Prokaryotic assemblies with no genes predicted by SEARCH_16S. Columns are: *Assembly*, Genbank accession; *Name*, scientific name, *Frag*. is Y if SEARCH_16S reports a fragment, otherwise N; *BLAST* gives the length and identity of the top BLAST hit to GG97, if any. In the first three assemblies (no shading), there is no BLAST hit with E<10, confirming that the assembly is missing the 16S gene. In the following five assemblies (light shaded), the BLAST hits confirm the presence of partial genes, indicating assembly errors and supporting predictions of fragments by SEARCH_16S. In the last four assemblies (dark shaded), the BLAST hit is consistent with a complete gene.

| Assembly | Name | Frag. | BLAST |
| --- | --- | --- | --- |
| GCA_000749465.2 | Amycolatopsis lurida NRRL 2430 | N | (no hit) |
| GCA_000988445.1 | Escherichia coli strain SQ171 | N | (no hit) |
| GCA_000714635.1 | Klebsiella pneumoniae KP5-1 | N | (no hit) |
| GCA_000019805.1 | Thermoproteus neutrophilus V24Sta | Y | 272 (100.0%) |
| GCA_001729625.1 | Bacterium AB1 | Y | 377 (100.0%) |
| GCA_000234805.1 | Pyrobaculum ferrireducens 1860 | Y | 402 (97%) |
| GCA_000281215.1 | Pseudomonas putida DOT-T1E | Y | 402 (100%) |
| GCA_000247545.1 | Pyrobaculum oguniense TE7 | Y | 467 (100%) |
| GCA_000016385.1 | Pyrobaculum arsenaticum DSM 13514 | Y | 1,340 (99%) |
| GCA_000007225.1 | Pyrobaculum aerophilum IM2 | Y | 1,521 (100%) |
| GCA_001189275.1 | Pyrobaculum sp. WP30 | Y | 2,126 (100%) |
| GCA_000011125.1 | Aeropyrum pernix K1 DNA | Y | 2,201 (100%) |

### Circular chromosomes containing split 16S genes

Many bacterial chromosomes are circular, and in these cases Genbank chromosome sequences start at arbitrary positions which may split a gene into two segments. SEARCH_16S therefore considers bases at the start of a chromosome to follow bases at the end. Split 16S genes were found in the following chromosomes: CP006046.2, CP006996.1, CP009571.1, CP010836.1, CP011479.1, CP013689.1, CP017279.1 and CP012986.1.

### Comparison with NCBI feature tables

Of the 6,487 finished assemblies, 6,401 had feature tables provided by NCBI. SEARCH_16S and NCBI reported the same number of 16S genes (i.e., number of paralogs) in 5,724 (88%). A total of 26,816 genes were reported with 24,402 (91%) reported by both. A gene was considered to be reported by both methods if the coordinates overlapped by at least 50%. I manually reviewed many of the annotations where the methods disagreed and found all of them to be problems in the feature tables. In some cases, the annotation was too long, possibly covering the complete SSU operon. I found 12 cases where the NCBI annotation had the correct coordinates on the wrong strand. 466 assemblies had no 16S genes in the feature tables but did have genes predicted by SEARCH_16S; these cases are clearly false negatives in the annotations rather than false positives by SEARCH_16S. These observations confirm previous reports (Lagesen *et al*., 2007; Freyhult *et al*., 2007) of unreliable 16S annotations in public databases.

### SILVA, RDP and Greengenes

Most sequences in SILVA, RDP and Greengenes are partial genes which lack the boundary motifs used by SEARCH_16S (see “Truncated sequences” below). In Greengenes, all sequences were reported as hits (genes or fragments). In the prokaryotic subset of SILVA, *99.99%* of sequences were hits. Of the 168 sequences with no hit, 157 were mitochondrial genes from eukaryotic cells which are classified as Proteobacteria by SILVA, and five were chloroplasts. The remaining six were all uncultured sequences. In the prokaryotic subset of RDP, only 84% of sequences were hits. This is due to the many partial genes in RDP which are too short to be reported as fragments (Fig. 5).

**Fig 5.**
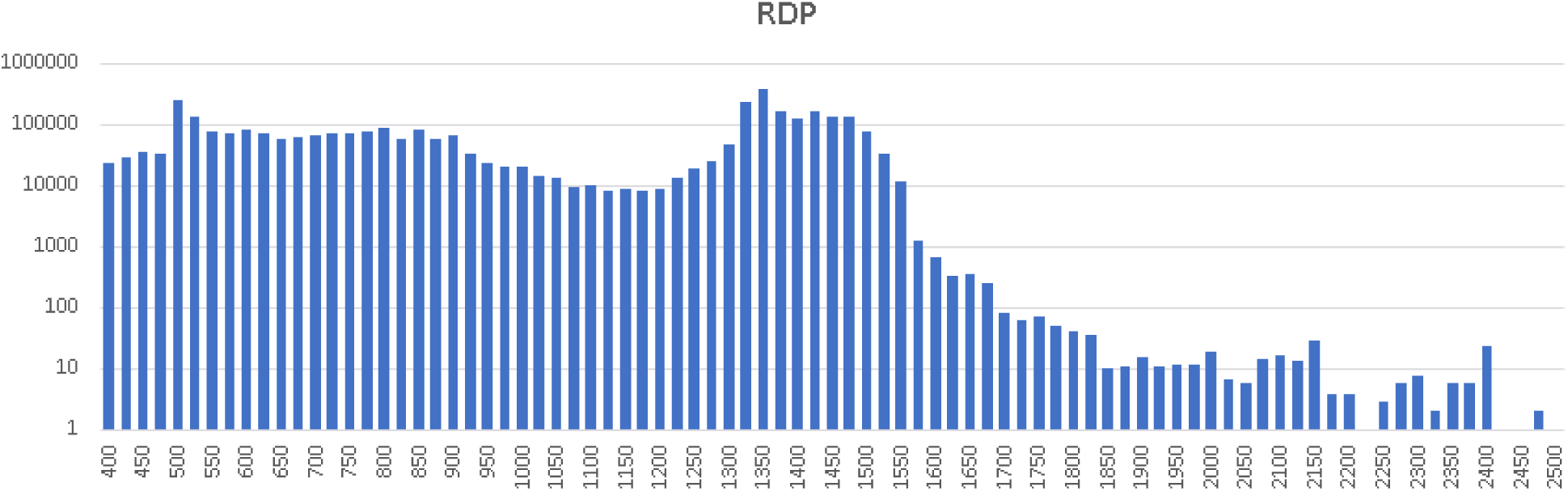
Length distribution of sequences in RDP. The histogram shows lengths binned into intervals of 25. Most sequences have lengths <1400nt and are therefore very likely to be partial genes (compare with Fig. 4). Many sequences have lengths <500nt and are therefore too short to be reported as fragments by SEARCH_16S.

### Truncated sequences in Greengenes and SILVA

I investigated the multiple alignments provided with Greengenes and SILVA and found that a large majority of sequences are missing bases at both ends. This was determined as follows. Almost all 16S genes in the finished assemblies contain bases which align to the C11F and C1512R motifs, with the exceptions summarized in Table 2. These regions are therefore rarely, if ever, deleted. I located the multiple alignment columns containing these motifs and measured the fraction of sequences with coverage in those columns, finding that a large majority of Greengenes and SILVA sequences are truncated after C11F and before C1512R (Fig. 6).

**Fig 6.**
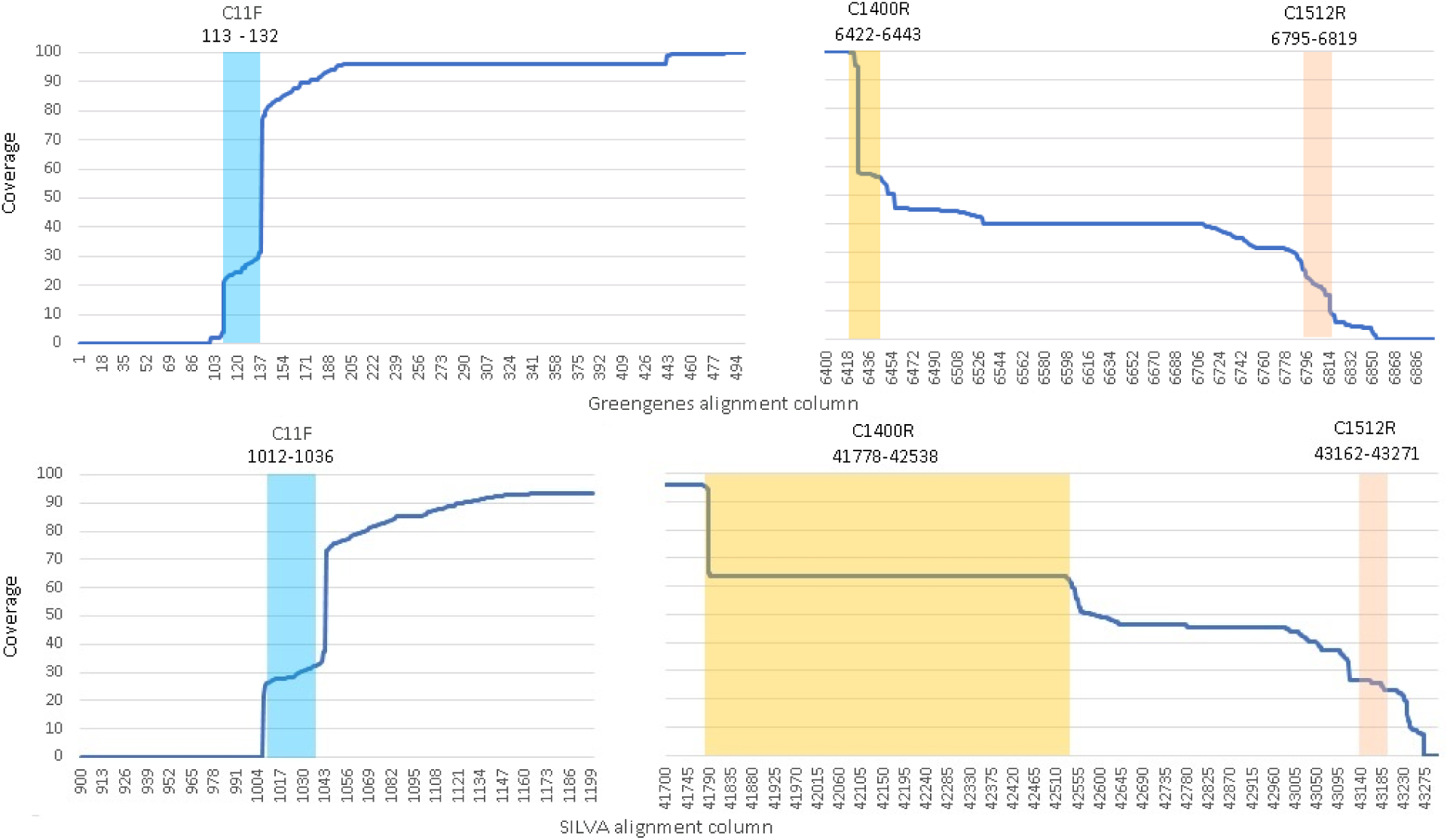
Truncated sequences in Greengenes and SILVA. Sequence coverage at the ends of the Greengenes (top) and SILVA (bottom) multiple alignments. A sequence is considered to cover a column if it has a letter or internal gap in that column, otherwise it contains a terminal gap and does not cover the column. This shows that many sequences are truncated such that they do not contain the C11F or C1512R motifs used by SEARCH_16S and will therefore be reported as fragments. Many sequences are truncated immediately before the C1400R motif (GGGTCTTGTACACACCG). We can therefore infer that many sequences were obtained by PCR with primers at, or close to, C11F at the start and C1400R or C1512R at the end.

### Eukaroytic subset of SILVA and metazoan subset of nt

In the eukaryotic subset of SILVA, 79 of the 120,702 sequences were hits (5 genes and 74 fragments). This is a hit rate of 0.07%, compared to 99.99% for the prokaryotic subset. On the metazoan subset of nt, SEARCH_16S reported 25 genes and 1,268 fragments in 12.3M sequences (0.01% hits). Two of the reported genes were annotated as 16S in Genbank (AY833572.1, *Bemisia tabaci* and JN975069.1, *Caenorhabditis elegans*). JN975069.1 is 98% identical to the 16S gene in NCBI reference sequence NR_074668.1 (*Burkholderia phymatum* strain STM815 16S ribosomal RNA gene) and is thus unambiguously a 16S gene. Six were mitochondria. Nine were from scaffolds in unfinished genomes, seven of which had identities >97% with GG97 and are therefore true positives by SEARCH_16S which are probably due to contaminants in the reads, or possibly to horizontal gene transfer from a prokaryote. Only one of these genes was annotated as 18S. Overall, these results show that SEARCH_16S is highly effective at distinguishing 16S from 18S.

### Random sequence

As a further test of specificity, I generated 10^5^ pseudo-random sequences of length 10^6^ letters using uniform probabilities for each nucleotide. SEARCH_16S reported no genes or fragments in these sequences.

### Comparison with RNAmmer

As noted in the Introduction, to the best of my knowledge RNAmmer is the only previously published specialized method for identifying 16S genes, and I therefore wanted to compare its predictions with SEARCH_16S on the same test datasets. Unfortunately, this was not possible as I was unable to obtain the software (I requested a license for validation purposes at http://www.cbs.dtu.dk/cgi-bin/sw_request?rnammer, but received no response). RNAmmer predictions on selected genomes are posted at http://www.cbs.dtu.dk/services/RNAmmer/. I downloaded these predictions on 15th Dec. 2016 and found that the files were dated 14th Jan. 2014 despite the statement on that page that it is updated daily. Only 978 genomes were included, far fewer than the 6,487 complete genome assemblies currently available. Genbank accession numbers are given without the version number, so for example there are annotations for AE000516 but it is not documented whether the sequence is AE000516.1 or AE000516.2, precluding a comparison with SEARCH_16S on the same set of sequences. I therefore chose to compare the number of 16S genes reported for each complete genome present in the downloaded RNAmmer predictions, reasoning that if the genome is annotated as “Complete”, then the number of 16S genes is unlikely to change if the sequence is updated. I found that RNAmmer and SEARCH_16S agreed on the number of genes in 966 / 978 (99%) of those genomes.

### CPU time and memory use

Execution time and memory use on the test datasets is summarized in Table 3. The tests were run using the search_16s command in the 64-bit Windows build of USEARCH v10.0 on an Intel i7 PC with 6 cores and 32 Gb of RAM.

**Table 3.** Execution time and memory use on the test datasets. The tests were run on an Intel i7 PC with 6 CPU cores.

| Dataset | File size | Time | RAM |
| --- | --- | --- | --- |
| Complete genomes | 24 Gb | 6m 45s | 1.3 Gb |
| Greengenes | 1.7 Gb | 50s | 129 Mb |
| RDP (prok.) | 3.9 Gb | 1m 40s | 120 Mb |
| SILVA (prok.) | 2.4 Gb | 1m 17s | 120 Mb |
| SILVA (euk.) | 216 Mb | 5s | 116 Mb |
| Metazoa | 42 Gb | 12m 25s | 2.1 Gb |
| Random | 100 Gb | 20m 10s | 120 Mb |

## Discussion

The SEARCH_16S algorithm is remarkably fast and accurate. If reports of genes and fragments are interpreted appropriately, the results reported here show that it is almost perfectly sensitive with no detectable errors. For genome annotation, genes are almost certainly correct and fragments should be examined manually to determine whether they are intact genes or assembly errors. 18S genes may be reported as fragments, or in very rare cases may be reported as genes.

## References

Acinas, S.G. et al. (2004) Divergence and redundancy of 16S rRNA sequences in genomes with multiple rrn operons. J. Bacteriol., 186, 2629–35.

Altschul, S.F. et al. (1997) Gapped BLAST and PSI-BLAST: A new generation of protein database search programs. Nucleic Acids Res., 25, 3389–3402.

DeSantis, T.Z. et al. (2006) Greengenes, a chimera-checked 16S rRNA gene database and workbench compatible with ARB. Appl. Environ. Microbiol., 72, 5069–72.

Eddy, S.R. (2009) A new generation of homology search tools based on probabilistic inference. Genome Informatics, 23, 205–211.

Freyhult, E.K. et al. (2007) Exploring genomic dark matter: A critical assessment of the performance of homology search methods on noncoding RNA. Genome Res., 17, 117–125.

Lagesen, K. et al. (2007) RNAmmer: Consistent and rapid annotation of ribosomal RNA genes. Nucleic Acids Res., 35, 3100–3108.

Maidak, B.L. et al. (2001) The RDP-II (Ribosomal Database Project). Nucleic Acids Res., 29, 173–4.

Pruesse, E. et al. (2007) SILVA: A comprehensive online resource for quality checked and aligned ribosomal RNA sequence data compatible with ARB. Nucleic Acids Res., 35, 7188–7196.

Sayers, E.W. et al. (2012) Database resources of the National Center for Biotechnology Information. Nucleic Acids Res., 40, D13–25.

Shajani, Z. et al. (2011) Assembly of Bacterial Ribosomes. Annu. Rev. Biochem., 80, 501–526.

